# Genetic variation associated with relative resistance in teak (*Tectona grandis* L. f.) against the leaf skeletonizer, *Eutectona machaeralis* Walker

**DOI:** 10.1101/2022.08.10.503439

**Authors:** Vivek Vaishnav, Nitin Kulkarni, Shamim Akhtar Ansari, Tikam Singh Rana

**Affiliations:** Molecular Systematic Laboratory, Plant Diversity, Systematics and Herbarium Division, CSIR- National Botanical Research Institute, Lucknow, 226001, India; Laboratory for Conservation and Genetic Improvement of Forest Trees Department of Forestry and Environmental Science Manipur University, Indo-Myanmar Road, Canchipur, Imphal, Manipur, 795003, India; Institute of Forest Productivity, Gumla Road, Ranchi, Jharkhand, 482021, India; 206, Gemini Apartment, Khurram Nagar, Lucknow, 226022, India

**Keywords:** Forest Entomology, Integrated Pest Management, Heterozygosity, Systematic Acquired Resistance, Expressed Sequence Tags

## Abstract

**Background:** Photosynthesizing tissue of teak (*Tectona grandis* L. f.) foliage is damaged by a host-specific insect pest called leaf skeletonizer (*Eutectona machaeralis* Walker) that severely eclipses annual growth increment and carbon sequestration of natural populations and plantation of teak. Gene-assisted selection of relatively resistant teak clones may efficiently control the damage in the populations and plantations. The present investigation aimed to identify genetic variation associated with relative resistance in teak against the pest.

**Method:** The investigation was carried out on 106 teak plus tree clones assembled at the National Teak Germplasm Bank from the Indian meta-population of teak. Resistance data were obtained recording the ocular damage caused by the pest to teak accessions for four years. Genotyping of the teak accessions was performed with 21 co-dominant markers and marker-trait association mapping was performed confirming the genetic structure of the germplasm bank and linkage disequilibrium (LD) among the marker loci.

**Results:** The sampled teak accessions exhibited a low albeit highly admixed genetic structure (F_ST_=0.07) and low level of LD (16.66%) among loci, making them suitable for high-resolution association analysis. A significant correlation (*p*≤0.01, *R*^2^=0.67) was obtained between intra-specific heterozygosity and the relative resistance against the pest. A marker locus CCoAMT-1 representing the enzyme *caffeoyl-CoA O-methyltransferase* of phenylpropanoid pathway was also found significantly (*p*≤0.05) associated with the relative resistance against the pest explaining 6.6% of the phenotypic variation (*R*^2^=0.066) through positive effect (0.57) on the trait.

**Conclusions:** The present work exhibited a significant correlation of intra-specific heterozygosity with relative resistance in teak against a pest. It is the first report on teak identifying genetic markers associated with relative resistance against the pest. The marker can be applied for the selection of resistant planting stock for breeding and commercial plantation.

Further investigation can be performed to understand the expression level polymorphism linked with the resistance applying next-generation sequencing approaches.

## Introduction

Teak (*Tectona grandis* L. f) is a well-known timber species of South-Asian countries. Due to high commercial demand and ease of cultivation, the species plantations have been widely established throughout the tropics. It is emerging as a valuable hardwood resource in about 70 countries around the world, attracting large investment from the private sector in Africa, Asia, and Latin America (FAO 2015). The global annual trade of teak hardwood was about 3% of the global timber trade (FAO 2015). *Eutectona machaeralis* Walker (Lepidoptera: Pyralidae) is a major host-specific pest of teak that infests during August to October and specifically develops towards the end of the growing season before normal leaf shedding (Nair 2001). It is an oligophagous pest that feeds on mesophyll tissues, leaving only the leaf veins hence called ‘skeletonizer’. The skeletonizer has been reported to damage as much as half of the total annual increment of teak plantations (FAO 2003). Different studies from India have also reported a loss of 3%-8% in the annual increment of teak plantations (Sangeetha and Arivudainamb 2012) and damage of 55% of teak seedlings in the forest nurseries (Kulkarni *et al*. 2004) due to the pest skeletonizer. The control of the teak skeletonizer can therefore lead to substantial economic gain due to the high value of teak timber, and the large area under commercial plantations. Although some methods based on silvicultural-cum-biological control (Beeson 1941), and chemical control (Basu-Chowdhury 1971) have been recommended as the pest control measure, no method as such is currently employed as a preventive method to control the chances of the pest infestation (Shukla and Joshi 2001). Biological and chemical methods, to control pest infestation in crops, need repetition at a specific interval. In the case of forest trees, those methods may be cost-intensive, and could also affect the environment adversely. Therefore, an integrated pest management (IPM) approach using resistant planting stock has been suggested to avoid the chances of pest infestation (Grossi-de-Sá *et al*. 2015; Kulkarni 2017). Now, biotechnological tools have also been employed for regulating functional genomics through transgenesis for the development of resistant elite plants and gene silencing for down-regulating the pests for their control (Zhang *et al*. 2017).

Plants exhibit a considerable multitude of natural variations in defense mechanisms shaped by the different selection pressures (Thompson 2005). The natural variation in plant traits, like trichome density (Kaplan *et al*. 2009), leaf lamina area, and cuticle thickness (Moles *et al*. 2011) have been found significantly associated with herbivore resistance (Carmona *et al*. 2011). Secondary metabolites are also known to play a significant role in defense against herbivores with the pleiotropic effect of genes associated with them (Carmona *et al*. 2011). The selection of genetically improved genotypes utilizes the natural genetic variation of the plants for the trait. However, very little of the natural variation of forest trees could be exploited for their genetic improvement. The wide array of genetic heterogeneity can be explored to screen out the natural resistant genotypes and variety (Broekgaarden *et al*. 2011) with help of biotechnological interventions.

Association studies, based on the DNA marker system, helps to develop the gene-specific markers associated with specific resistance to facilitate marker-assisted selection (MAS) and breeding (Butcher and Southerton 2007). Numerous molecular markers developed in many tree species have enabled the genetic dissection of important quantitative characters using an association mapping approach. Efforts have been made to screen out the DNA markers linked with genes responsible for the resistance against pests in some model plant species (Broekgaarden *et al*. 2011; Sandhu and Kang 2017). Mapping wheat genome through microsatellite markers, two QTLs from 12-oxo-phytodienoic acid reductase (OPR) and lipoxygenase (LOX) genes was reported significantly associated with resistance to Hessian fly (*Mayetiola destructor*) in wheat (Tan *et al*. 2013). Through the QTL mapping approach, single nucleotide polymorphism (SNP) linked with QTL governing the resistance to sunn pests (*Eurygaster integriceps* Puton) was identified in wheat (Emebiri *et al*. 2017). In rice also, SNPs associated with resistance against brown planthopper (*Nilaparvata lugens* Stål) were reported (Kusumawati *et al*. 2018). These approaches have not been applied to non-model forest tree species due to a lack of background genomic information and the molecular level physiological mechanism regulating the resistance. There are only a few investigations reported for disease resistance (Quesada *et al*. 2010) and pest resistance (Zhang *et al*. 2018) in forest trees. Since teak is highly sought-after timber in the global market, we applied the association mapping approach to identify naturally introgressive genomic loci associated with the relative resistance in teak against the skeletonizer *E. machaeralis*.

## Methods

### Assessment of *E. machaeralis* infestation

National Teak Germplasm Bank (NTGB), situated at Chandrapur, Maharashtra, India (*N 19*.*976240 E 079*.*338117*), had been established planting triplicate ramets of plus trees (phenotypically superior trees) selected in natural forests, distributed over 12 states in India (Kumar *et al*. 1998). The relative resistance data against the infestation of *E. machaeralis* was observed on three ramets of 106 teak plus tree clones representing 10 states (Roychoudhury and Joshi 1996). The continuous variation in relative resistance among the genotypes against the pest infestation was recorded based on ocular observation of defoliation in categories of 0%, ≤25%, ≤50%, ≤75%, and ≤100%, respectively for every year in each ramet (Additional Table 1). For investigation, these categories were assessed through five rating criteria, viz., 0, 1, 2, 3, and 4 respectively (Additional Table 1). An average rating of four years of observation for each ramet was obtained in nine different classes (Roychoudhury and Joshi 1996) covering the most susceptible (3.0) to the most resistant (1.0) that was further employed for the genetic analysis (Table 1).

**Table 1.** Genetic heterogeneity of the teak among nine classes of rating for resistance, covering relatively the most susceptible (3.0) to the most resistant (1.0) genotypes against teak skeletonizer

| Mean Rating | N | %P | Ho |
| --- | --- | --- | --- |
| 3.00 | 2 | 57.14 | 0.09 |
| 2.75 | 5 | 71.43 | 0.21 |
| 2.50 | 10 | 100.00 | 0.15 |
| 2.25 | 20 | 100.00 | 0.19 |
| 2.00 | 14 | 100.00 | 0.15 |
| 1.75 | 24 | 100.00 | 0.22 |
| 1.50 | 21 | 100.00 | 0.25 |
| 1.25 | 9 | 95.24 | 0.27 |
| 1.00 | 1 | 47.62 | 0.47 |

N- Number of genotypes, %P- percentage of polymorphism, Ho- observed heterozygosity

### Plant materials for DNA isolation

The branch cuttings collected from each of 106 teak tree clones were sampled from the NTGB. The sampled cuttings were treated with IAA (5 mM) at the bottom end and sealed at the top end with the paraffin wax, and then planted in the nursery of Genetics and Plant Propagation Division, Tropical Forest Research Institute, Jabalpur. The newly sprouted juvenile leaves from those cuttings were harvested and stored in a cryo-freezer (-80 °C) for genomic DNA isolation.

### DNA isolation and marker assay

The DNA was extracted from the leaf samples following a modified CTAB method (Narayanan *et al*. 2006). A set of 21 DNA markers including 13 nuclear simple sequence repeats (nuSSRs) primers and 8 expressed sequence tag (EST) based markers (Vaishnav and Ansari 2018) were employed for the investigation. The EST-based markers included a Catalase gene-based primer characterized in teak (Cheua-ngam and Volkaert 2006). To employ candidate gene-based approach in association analysis, 15 primers were designed from the genes representing three enzymes of the phenylpropanoid pathway in plants *viz. phenylalanine ammonia-lyase* (PAL), *cinnamoyl-CoA reductase* (CCR), and c*affeoyl-CoA O-methyltransferase* (CCoAMT), following the primer designing criteria and procedure as described by Vaishnav and Ansari (2018). Out of these 15 primers, only 7 primers could be characterized on the sampled teak genotypes (Table 2). Finally, 21 sets of primers were selected for the final amplification (Additional file, Table 2). The PCR amplification was performed following the protocol as described by Vaishnav and Ansari (2018). A data profile was generated from the electrophoresis gel images. The samples with missing alleles (<5%) were omitted and finally, a genetic profile of 21 co-dominant markers on 106 teak genotypes along with their corresponding relative resistance data was developed for analysis.

**Table 2.**
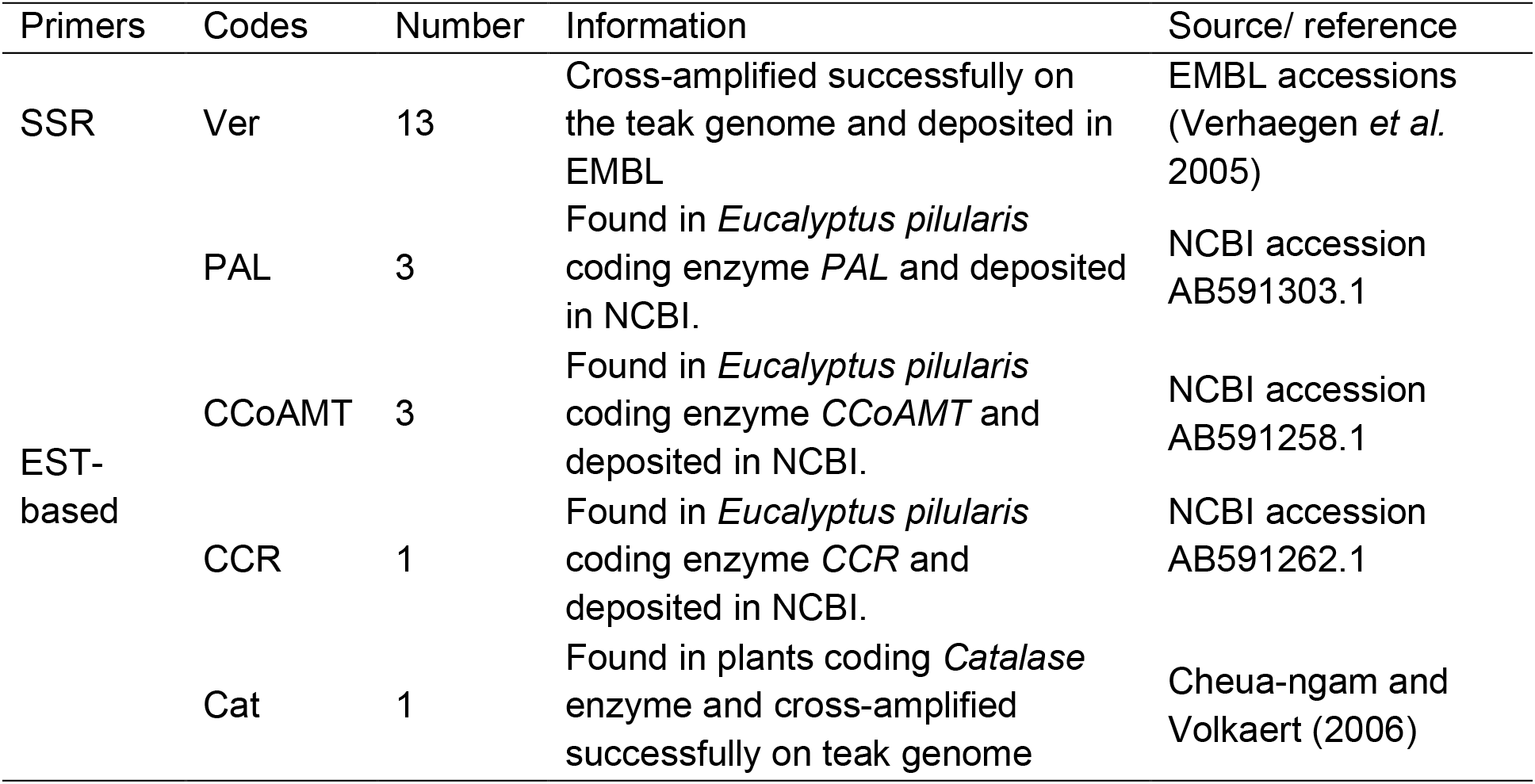
Information of microsatellites designed and adopted for cross-amplification on teak (*Tectona grandis* L.f.)

### The genetic information of the markers

Major allele frequency (*MAF*), expected heterozygosity or gene diversity (*H*_*e*_), observed heterozygosity (*H*_*o*_), and the polymorphic information content (*PIC*) were calculated for each marker using POWERMARKER v3.25 (Liu and Muse 2005). To identify the loci with a signature of environmental adaptation, a highly robust program BAYESCAN v2.1 (Foll and Gaggiotti 2008) was applied to detect the loci under selection. The q value threshold, i. e. ≤ 0.1 was used to discriminate false positive among the detected loci employing reversible jump MCMC (burn-in 50 000 iterations, a thinning interval 20, and a sample size of 5000) and estimation of the Bayesian posterior probability in form of logarithm value of the posterior odds (Log_10_ (PO).

### Genetic structure and linkage disequilibrium (LD)

Genetic diversity and structure were estimated to confirm true representation of the natural variation in the teak genetic resource of the country by the germplasm bank, and to avoid the chances of spurious and false-discovery of association due to a structured sample. A Bayesian analysis of genetic structure was performed using STRUCTURE v2.3.4 (Pritchard *et al*. 2000; Falush *et al*. 2007; Hubisz *et al*. 2009). The analysis uses multi-locus genotypes to infer the fraction of an accession’s genetic ancestry that belonged to a population, for a given number of populations (K). The posterior probabilities were estimated for each value of K between 1 and 10 with MCMC simulation. The results were based on 500,000 iterations of this chain, following a burn-in period of 1000,000 iterations. The MCMC chain was run multiple times, using LOCPRIOR with admixture and correlated allele frequency model (prior mean is 0.01, prior SD 0.05, and lambda set at 1.0 in the advanced option of the program). For every number of K three simultaneous runs were performed. To infer the true number of K, the ΔK method developed by Evanno *et al*. (2005) was implemented with the help of online software STRUCTURE HARVESTER (Earl and VonHoldt 2012). Following the most appropriate number of K assigned for the samples resulted in the software, the corresponding inferred ancestry coefficient (Q) for each genotype was obtained. A model-free approach was also applied to discriminate the genetic structure of investigated teak accessions and principal coordinate analysis (PCoA) was performed through program GENALEX v6.0 (Peakall and Smouse 2012). An analysis of molecular variance (AMOVA) was performed in ARLEQUIN v3.5 (Excoffier and Lischer 2010) to obtain the variation in genetic differentiation among representative population accessions and their F_ST_ was also calculated. The level of LD among loci was estimated using the pair-wise recombination coefficient (R^2^ values) calculated by TASSEL v2.1 (Bradbury *et al*. 2007). The percentage of the marker combinations was calculated by estimating the number of marker combinations with significant LD (p≤0.05) on the total number of marker combinations at a different level of R^2^ values >0.1.

### Genetic heterogeneity of relative resistance

Genetic heterogeneity of teak genotypes among nine classes covering relatively the most susceptible (3.0) to the most resistant (1.0) against the skeletonizer was estimated by measuring the percentage of polymorphism (*% P*), and *H*_*o*_ applying program POPGENE v1.31 (Yeh *et al*. 1999).

### Marker-resistance association and verification

The marker-trait association analysis was performed applying both general and mixed linear models (GLM and MLM) utilizing the algorithm efficient mixed-model association or EMMA (Kang *et al*. 2008) through TASSEL v2.1 (Bradbury *et al*. 2007), as suggested by Yu *et al*. (2006) incorporating a kinship matrix (K; calculated through TASSEL) and the ancestry coefficient (Q; obtained from STRUCTURE) to control false-discovery rate (FDR). Advanced over GLM, the MLM considers the markers applied to the study as a fixed–effect factor, and the population structure information of the sampled genotypes are considered as random effect factors avoiding possible spurious association. The significant marker-trait association was determined based on marker adjacent *p*-value (*p*≤0.05). Bonferroni correction was applied (*α*=0.05) to screen out the false-discovery of association. The *R*^*2*^ value indicated the percentage of phenotypic variance explained by the identified marker associated with the trait. The phenotypic effect of the allele was also determined and the mean of positive or negative allele effects was calculated as the average (positive or negative) allelic effect of a marker/locus.

## Results

### The genetic information of the markers

The analysis of the genetic profile of teak plus tree clones on 21 co-dominant markers resulted in a major allele frequency of 0.67±0.11 ranging from 0.52 to 0.90 by the markers. The *H*_*e*_ was 0.41±0.09 ranging from 0.18 to 0.50, whereas *H*_*o*_ was 0.21±0.20 that varied from 0 to 0.75. The *PIC* value was 0.32±0.05, which ranged from 0.16 to 0.37 (Additional Table 2). The F_ST_ values of the markers ranged from 0.11 to 0.19 and no locus was found as an outlier under selection due to insignificant (<0.5) posterior odd (Figure 1).

**Figure 1.**
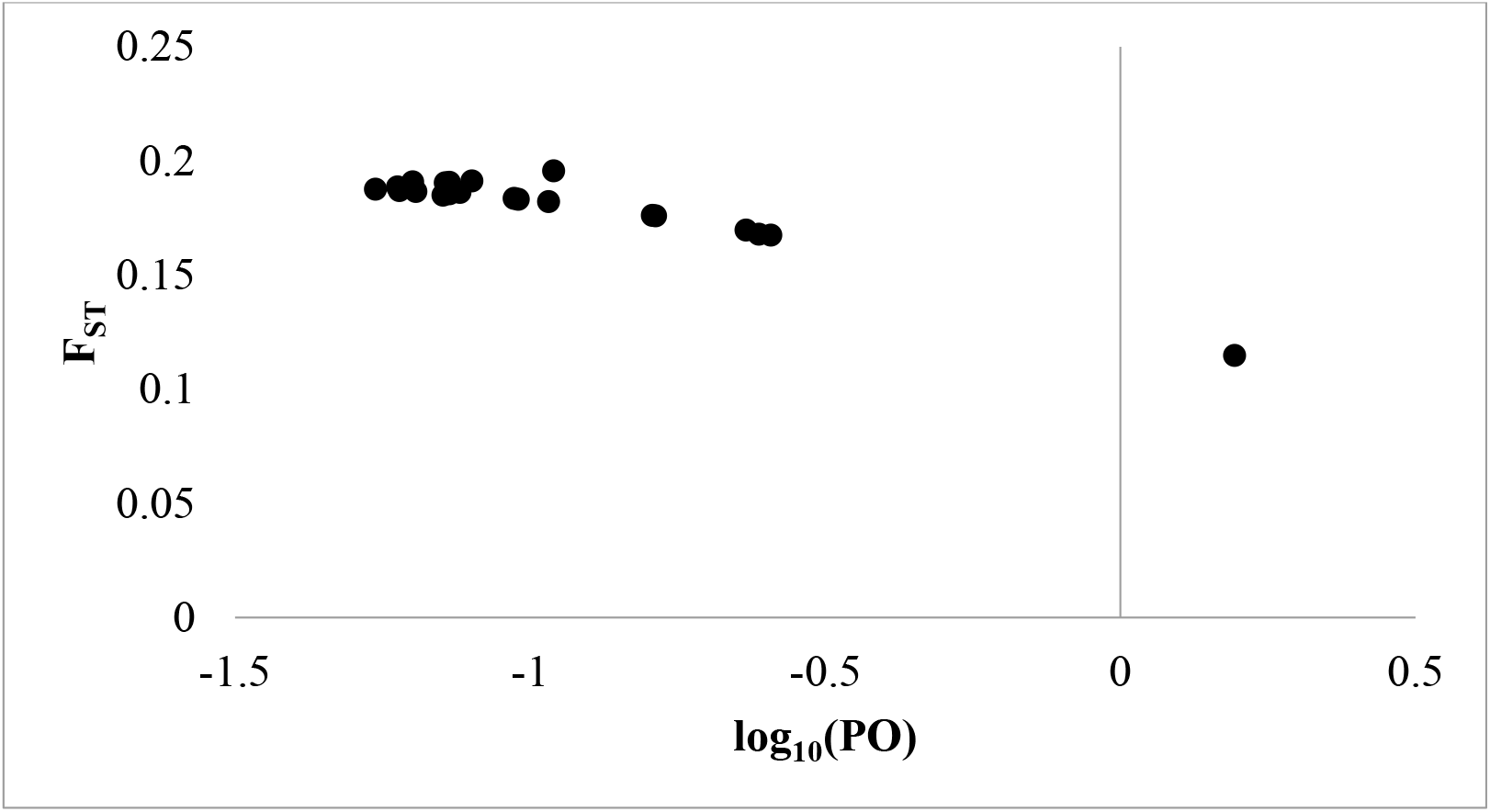
Bayescan analysis resulting F_ST_ values on Bayes Factor, i.e. Log_10_ (PO) for 21 markers, confirms no outlier (Log_10_ (PO) >0.5) depicting no evidence of selection.

### Genetic structure and linkage disequilibrium

Teak plus tree clones sampled for the investigation exhibited 65.24±34.70% of polymorphism, and 0.29±0.15 of *H*_*e*,_ and 0.27±0.14 of observed heterozygosity, respectively (Additional Table 3). The Bayesian estimation-based genetic structure assessment of the germplasm bank resulted in very low genetic structure (ΔK<250) with low support to K=2 as the most suitable number of cryptic populations for the sampled teak genotypes (Additional file, Figure 1). The PCoA also resulted in one admixed group of six locations of teak from the north, central and south India and the other four locations of the east (ARP), west (GJ), central (MP), and south (KR) India remained distinctly (Additional Figure 2). The hierarchical variation was 7.48% among the representative populations and 92.51% within the population with an F_ST_ value of 0.07. Among the 210 pairs of loci, 16.66% were found in significant LD (*p*≤0.05, *R*^*2*^ values > 0.1).

**Figure 2.**
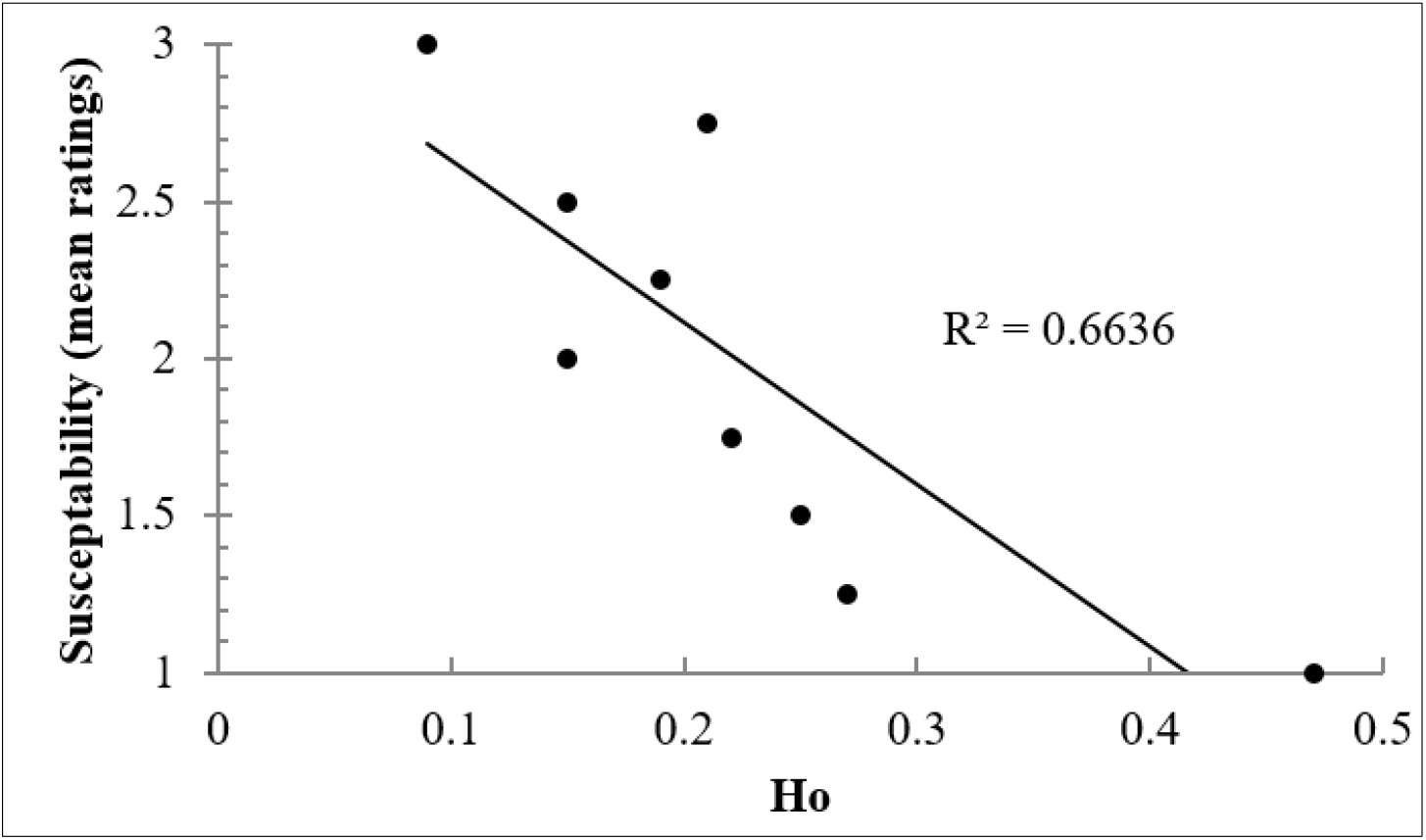
Observed heterozygosity (*H*_*o*_) of teak genotypes in different relative resistance classes was observed declining from high resistant (1.0) to high susceptible (3.0) classes of teak against teak skeletonizer

### Genetic heterogeneity of relative resistance

The polymorphism in different classes of the relative resistance of the representative teak trees ranged from 47.62% to 100% while *H*_*o*_ varied from 0.09 to 0.47 (Table 1). A highly significant (*p*≤0.01) and positive correlation was found between relative resistance class and intra-specific *H*_*o*_ (Figure 2).

### Genomic loci associated with the relative resistance

Among all markers, only the CCoAMT-1 was found significantly (*p*≤0.05) associated with the relative resistance to the skeletonizer in teak explaining 6.6% of the phenotypic variation (*R*^*2*^= 0.066) through positive effect (0.57) on the trait.

## Discussion

Teak is an entomophilous tree species with a high crossing rate (Shrestha *et al*. 2005), hence insects play a major role in pollination (gene flow) in teak meta-population maintaining a greater variation among the genotypes within the population (Verhaegan *et al*. 2010; Vaishnav *et al*. 2015; Vaishnav and Ansari 2018). Application of chemical pesticides against the pests may result in the mortality of the useful insects helping its pollination success and genetic diversity. A very high intra-specific genetic diversity exhibited by teak populations needs high selection intensity for increased genetic gain for quantitative traits such as relative resistance against the pest. Therefore, there is a wide scope for the DNA-based marker-assisted selection of resistant teak genotypes. But lack of investigation on genetic causal factors governing the resistance against pests like teak skeletonizer has led to a gap in molecular level information to apply the marker-assisted selection of resistant planting material.

The advancement of gene sequencing technology and available genic sequences in genome database (*e. g*. National Center for Biotechnology Information) has improved the robustness and resolution of marker-trait association analysis. Therefore, we preferred a candidate gene-based approach of association mapping and applied few co-dominant markers from the gene sequences representing enzymes of the phenylpropanoid pathway, discovered in model plant species. The phenylpropanoid pathway producing the lignin and other secondary metabolites are known to have a major role to express the resistance in different plant species to various pests and pathogens (Raiskila 2008; Santiago *et al*. 2013). Enzymes of this pathway were found regulating the synthesis of secondary metabolites providing structural support to the plant wood and inducing the systematic acquired resistance (SAR) against pests and pathogens (Sticher *et al*. 1997; Fraser and Chapple 2011). *Phenylalanine ammonia-lyase* (PAL), the regulating enzyme of the phenylpropanoid pathway, converts phenylalanine into cinnamate. It has been found associated with resistance in plants (Dixon *et al*. 2002; Ning and Wang 2018). *Cinnamoyl-CoA reductase* (CCR), which converts p-coumaroyl Co-A into p-coumaraldehide, is found to have a role in defense signaling in plants (Kawasaki *et al*. 2006). *Caffeoyl-CoA O-methyltransferase* (CCoAMT) converts Caffeoyl Co-A into Feruloyl Co-A and has been known to trigger disease resistance response in plants (Schmitt *et al*. 1991). Since these enzymes have been discovered to play role in the regulation of pest resistance in plants, we preferred them to design primers for our investigation.

The collection of NTGB was investigated as a training population for association analysis, as it is the only collection of the teak accessions in India that represents major provenances of teak metapopulation in form of a clonal orchard and suitable to perform a common garden test. Further, the germplasm bank collection is advantageous over natural population for marker-trait association studies, as in contrary to the samples from a germplasm bank collection, the phenotypic variation in the samples from a different natural population are critically influenced by the environment (Ingvarsson and Street 2011).

### The genetic information of the markers

The SSRs and ESTs have been successfully applied for the association mapping in many out-crossing plant species to identify the genomic loci linked to several traits of interest (Motilal *et al*. 2016; Zhang *et al*. 2016; Zhang *et al*. 2018, Vaishnav and Ansari 2018). For the present investigation, we applied 13 SSR markers and 8 EST-SSR markers specifically designed and characterized on teak. The genetic informativeness of these markers has been confirmed for high-resolution association analysis. The genetic parameters depicting the genetic information did not differ markedly among the markers (Additional Table 2). The major allele frequency (0.67±0.11) infers the evolutionary demographic history of a species genome (Kim *et al*. 2011) hence the value confirms the wide genome coverage of the markers for association mapping. The difference between the *H*_*e*_ and the *H*_*o*_ of the markers confirmed their deviation from Hardy-Weinberg Equilibrium (HWE) and two markers (Ver12 and Ver13) revealed slightly higher values of *H*_*o*_ than *H*_*e*_. Possibly, these two markers belong to the genomic region of the species that could be swiped in due to selected teak plus trees for the germplasm bank. In general, artificial selection based on superior traits leads to the heterozygote advantage (Hedrick, 2015). In the present study, the markers found with lower *H*_*o*_ than *H*_*e*_ confirmed the scope for a further selection of superior clones among the assembled accessions. Due to the specificity, the PIC values of the markers were found lower than those reported in other plant species (Kesari *et al*. 2010). No locus was found potentially affected by the divergent selection (Figure 1) avoiding historical demographic influence on genetic structure and differentiation of the germplasm (e.g. Mandel *et al*. 2013) hence could be reliable to apply for the marker-trait association analysis.

### Genetic structure and linkage disequilibrium

The teak population of India harbors the greatest genetic variability, revealing 80.30% to 80.55% of polymorphism and 0.64 to 0.76 of *H*_*e*_ (Fofana *et al*. 2008). However, the sampled genotypes with low values for polymorphism and *H*_*e*_ indicates the signature of genetic drift conceivably due to assemblage of only selected superior clones in the teak germplasm bank. The Bayesian-algorithm based analysis has revealed a very weak (ΔK <250) genetic structure of two cryptic populations (i.e. K=2), vis-a-vis high admixing among sampled genotypes. The PCoA has also presented similar results, where the source locations (states) of the teak genotypes have been found in the admixed group and have not presented a clear structure among them. Low F_ST_ (0.07) value confirms a high gene flow among sub-populations on an evolutionary time frame that has led to highly admixed Indian teak meta-population since the distant past. Due to short-distance cross-pollination through insects in teak (Shrestha *et al*. 2005), AMOVA has resulted in a higher genetic variation among the genotypes (92.51%) and avoids inbreeding depression in the population. Consequently, the observed significantly low LD (16.66%) among loci-pairs in the present investigation is feasible as allogamy attracts faster LD decay than autogamy. Hence it confirmed the suitability of the germplasm bank assemblage as a training population to exploit the historical recombination events for a fine resolution marker-trait association mapping (Myles *et al*. 2009). On the other hand, it suggests validating the association in a testing population to avoid the probability of false discovery.

### Genetic heterogeneity of relative resistance

A significant linear correlation between the relative resistance classes representing the most resistant (1.0) to the most susceptible (3.0) teak genotypes with observed heterozygosity (Figure 2) establishes that the intra-specific heterozygosity in the teak population may lead to the greater resistance against teak skeletonizer infestation. Although heterozygosity has been considered to contribute to population fitness against the herbivore (Egan and Ott, 2007), the investigation establishing intra-specific heterozygosity for pest resistance is rare. Mopper *et al*. (1991) found heterozygous trees of Pinus edulis resistant to the herbivores and concluded that the higher heterozygosity might have contributed to resistance against the pest. Moreover, teak is known for its durable timber quality that is found resistant against termites as well (Tewari, 1999). Such intra-specific variation for the pest resistance might have resulted due to mutation or evolutionary recombination history of polygene responsible for the relative resistance of teak against the continuous seasonal infestation of teak skeletonizer. However, a species-specific investigation reported that the plant genetic diversity effects on predators were independent of herbivores (Koricheva and Hayes 2018), therefore, a thorough investigation can be conducted in the future to validate the positive effect of intra-specific heterozygosity on pest resistance in plants.

### Genomic loci associated with the relative resistance

The enzyme CCoAMT catalyzes the methylation of caffeoyl-CoA in the phenylpropanoid pathway, which leads from trans-4-coumaroyl-CoA to trans-feruloyl-CoA (Matern and Kneusel 1988; ZH *et al*. 2001). Under any pest or pathogen attacks, the extent of change in the cell wall determines the course of infection depending on the reaction of the above pathway for resistance to the pathogens (Liu *et al*. 2018). As a response, feruloyl-CoA plays a major role in esterification for cell wall polysaccharides and lignifications making the key enzyme CCoAMT hyper essential for resistance. In an early investigation of its kind, the CCoAMT related gene ZmCCoAOMT2 is associated with resistance to multiple pathogens in Zea mays (Yang *et al*. 2017). Now in our investigation also, the locus CCoAMT-1 exhibited a significant association with the relative resistance against the pest teak skeletonizer. Since the function of the enzyme in the phenylpropanoid pathway for biosynthesis of monolignols has already been known, its association with resistance to teak skeletonizer strengthens the earlier findings regarding the role of lignin in plant resistance. We found the locus CCoAMT-1 contributed only a small fraction (6.6%) of resistance against the pest. On the replication of our finding of the association of enzyme involved in lignin biosynthesis with pest resistance, a number of QTLs can be developed related to the enzymes involved in the pathway, and those can indirectly be employed to screen out the elite teak genotypes with pest resistance. The quantitative inheritance of resistance trait fixes multiple loci with the relative and small effect of every locus during the polygenic inheritance. Therefore, the advanced molecular mapping techniques may help to localize and organize the other genes involved in pest resistance in teak.

## Conclusion

Genetic improvement of teak in India recommends ‘local source planting’ for the restoration of natural teak populations in India. A significant positive correlation between intra-specific heterozygosity and relative resistance advocates selection of heterozygote superior or development of hybrid vigor clones of teak with resistance against skeletonizer to avoid the chances of pest infestation and for the restoration of its natural populations. It is rare to achieve complete resistance or immunity against pests, as it may differ depending on spatial and temporal variables. Therefore, most of the tree improvement programs aim to achieve relative resistance to reduce the damage by the pests to a tolerable level. Identification of genetic causal factors or genes related to the plant resistance has always been given importance over other measures adopted for pest control that may cause harm to the beneficial insects which help in pollination and gene flow among the host plants. Therefore, the MAS would be of great importance to single out the pest-resistant host plants for breeding, conservation, and plantation program. The locus CCoAMT-1 related to the *CCoAMT* enzyme can be employed for genomic selection of pest-resistant teak planting stocks, after validation of our finding in a test population. Nevertheless, considering the quantitative nature and continuous variation in the pest-resistance along with the small genome coverage of applied markers, further association mapping should be investigated employing a large number of markers to understand the expression level polymorphism linked with the resistance applying next-generation sequencing with quantifying the relative expression-based analysis.

## List of abbreviations

(OPR): 12-oxo-phytodienoic acid reductase
(AMOVA): Analysis of molecular variance
(CCoAMT): Caffeoyl-CoA O-methyltransferase
(CCR): Cinnamoyl-CoA reductase
(He): Expected heterozygosity
(EST): Expressed sequence tag
(IPM): Integrated pest management
(LD): Linkage disequilibrium
(LOX): Lipoxygenase
(MAF): Major allele frequency
(MAS): Marker-assisted selection
(NTGB): National Teak Germplasm Bank
(nuSSRs): Nuclear simple sequence repeats
(Ho): Observed heterozygosity
(PAL): Phenylalanine ammonia-lyase
(PIC): Polymorphic information content
(PCoA): Principle co-ordinate analysis
(QTL/QTLs): Quantitative trait locus/loci
(SNP): Single nucleotide polymorphism

## Authors’ contributions

VV performed the field sampling and laboratory work, analyzed the data and prepared the first draft of the manuscript under the supervision of SAA and NK. SAA supported interpreting the molecular genetic part and NK supported in the entomology part of the investigation verifying the data and analysis.TSR drafted the final version of the manuscript through critical review for submission. All authors read and approved the final manuscript.

## Acknowledgements

Institutional support from Director, Institute of forest productivity, Ranchi, India and Director, Tropical Forest Research Institute, Jabalpur, India is gratefully acknowledged. Authors are grateful also to Director, Maharashtra Van Sansodhan Sansthan, Chandrapur, to provide access to the National teak germplasm bank. Fellowship support from the Council of Scientific and Industrial Research (CSIR), India to ‘VV’ is acknowledged.

