## Additional Tables and Figures for "Genetic variation associated with relative resistance in teak (*Tectona grandis* L. f.) against the leaf skeletonizer, *Eutectona machaeralis* Walker"

**Additional Table 1** Data-sheet showing ocular observation of damage caused by leaf skeletonizer *E. macheralis* in teak plus tree clones, recorded for consecutive four years. A five rating criteria, *viz.*, 0, 1, 2, 3 and 4 for was used to depict the defoliation of 0%, ≤25%, ≤50%, ≤75% and ≤100%, respectively

| SN | Accessions | Year-wise observation | | | | Average | R_Class |
|---|---|---|---|---|---|---|---|
| | | 1988 | 1989 | 1990 | 1991 | | |
| 1 | AP_58 | 2 | 2 | 2 | 2 | 2 | 5 |
| 2 | AP_60 | 2 | 2 | 2 | 1 | 1.75 | 6 |
| 3 | APJNB_1 | 1 | 1 | 2 | 2 | 1.5 | 7 |
| 4 | APKEA_23 | 1 | 1 | 2 | 2 | 1.5 | 7 |
| 5 | APKEA_24 | 1 | 1 | 3 | 3 | 2 | 5 |
| 6 | APKEA_25 | 2 | 1 | 2 | 2 | 1.75 | 6 |
| 7 | APKEC_1 | 1 | 1 | 2 | 3 | 1.75 | 6 |
| 8 | APKEC_2 | 1 | 2 | 2 | 2 | 1.75 | 6 |
| 9 | APKEN_I | 1 | 2 | 3 | 1 | 1.75 | 6 |
| 10 | APMN_4 | 2 | 1 | 2 | 4 | 2.25 | 4 |
| 11 | APNLP_1 | 2 | 3 | 2 | 2 | 2.25 | 4 |
| 12 | APNPL_11 | 1 | 2 | 3 | 3 | 2.25 | 4 |
| 13 | APNPL_2 | 1 | 1 | 1 | 2 | 1.25 | 8 |
| 14 | APNPL_5 | 2 | 2 | 2 | 1 | 1.75 | 6 |
| 15 | APNPL_6 | 1 | 2 | 2 | 1 | 1.5 | 7 |
| 16 | APNPL_7 | 2 | 1 | 2 | 1 | 1.5 | 7 |
| 17 | APNPL_8 | 2 | 2 | 2 | 1 | 1.75 | 6 |
| 18 | APT_10 | 3 | 2 | 2 | 3 | 2.5 | 3 |
| 19 | APT_11 | 3 | 2 | 2 | 3 | 2.5 | 3 |
| 20 | APT_14 | 2 | 3 | 3 | 2 | 2.5 | 3 |
| 21 | APT_15 | 2 | 1 | 2 | 4 | 2.25 | 4 |
| 22 | APT_16 | 1 | 1 | 2 | 3 | 1.75 | 6 |
| 23 | APT_17 | 2 | 1 | 2 | 2 | 1.75 | 6 |
| 24 | APT_20 | 1 | 1 | 2 | 1 | 1.25 | 8 |
| 25 | APT_22 | 3 | 2 | 3 | 2 | 2.5 | 3 |
| 26 | APT_3 | 2 | 2 | 2 | 3 | 2.25 | 4 |
| 27 | APT_6 | 1 | 2 | 2 | 3 | 2 | 5 |
| 28 | APT_7 | 3 | 2 | 2 | 3 | 2.5 | 3 |
| 29 | APT_8 | 3 | 2 | 2 | 2 | 2.25 | 4 |
| 30 | APT_9 | 2 | 1 | 2 | 3 | 2 | 5 |
| 31 | SBL_1 | 2 | 2 | 2 | 3 | 2.25 | 4 |
| 32 | AC_II | 1 | 2 | 2 | 4 | 2.25 | 4 |
| 33 | G+10 | 1 | 3 | 3 | 4 | 2.75 | 2 |
| 34 | KLN_2 | 2 | 1 | 2 | 3 | 2 | 5 |
| 35 | KLN_4 | 2 | 1 | 2 | 3 | 2 | 5 |

| | | | | | | | |
|---|---|---|---|---|---|---|---|
| 36 | KLS_2 | 2 | 2 | 2 | 4 | 2.5 | 3 |
| 37 | KLS_4 | 3 | 1 | 2 | 4 | 2.5 | 3 |
| 38 | MYHD_2 | 1 | 1 | 3 | 4 | 2.25 | 4 |
| 39 | MYHD_3 | 1 | 1 | 2 | 2 | 1.5 | 7 |
| 40 | MYHD_4 | 1 | 1 | 2 | 2 | 1.5 | 7 |
| 41 | MYHV_3 | 1 | 1 | 2 | 1 | 1.25 | 8 |
| 42 | MYHV_6 | 1 | 2 | 2 | 1 | 1.5 | 7 |
| 43 | MYSA_2 | 1 | 2 | 2 | 3 | 2 | 5 |
| 44 | ST_11 | 1 | 1 | 2 | 2 | 1.5 | 7 |
| 45 | ST_15 | 1 | 1 | 2 | 2 | 1.5 | 7 |
| 46 | ST_16 | 1 | 2 | 2 | 2 | 1.75 | 6 |
| 47 | ST_17 | 1 | 1 | 1 | 2 | 1.25 | 8 |
| 48 | ST_19 | 2 | 1 | 2 | 3 | 2 | 5 |
| 49 | ST_20 | 1 | 1 | 2 | 2 | 1.5 | 7 |
| 50 | ST_22 | 2 | 1 | 2 | 1 | 1.5 | 7 |
| 51 | ST_29 | 1 | 2 | 2 | 2 | 1.75 | 6 |
| 52 | ST_47 | 1 | 3 | 3 | 2 | 2.25 | 4 |
| 53 | ST_6 | 1 | 1 | 2 | 1 | 1.25 | 8 |
| 54 | ST_8 | 1 | 2 | 2 | 2 | 1.75 | 6 |
| 55 | MHAL_A2 | 2 | 1 | 3 | 1 | 1.75 | 6 |
| 56 | MHAL_A3 | 1 | 2 | 2 | 4 | 2.25 | 4 |
| 57 | MHAL_A4 | 2 | 1 | 2 | 2 | 1.75 | 6 |
| 58 | MHAL_A5 | 1 | 1 | 2 | 1 | 1.25 | 7 |
| 59 | MHAL_A6 | 2 | 2 | 2 | 2 | 2 | 5 |
| 60 | MHAL_A7 | 2 | 2 | 2 | 3 | 2.25 | 4 |
| 61 | MHAL_A8 | 2 | 2 | 2 | 3 | 2.25 | 4 |
| 62 | MHAL_A9 | 1 | 1 | 2 | 1 | 1.25 | 6 |
| 63 | MHAL_AI | 1 | 2 | 2 | 4 | 2.25 | 4 |
| 64 | MHAL_P2 | 3 | 2 | 2 | 4 | 2.75 | 2 |
| 65 | MHAL_P5 | 1 | 2 | 2 | 0 | 1.25 | 8 |
| 66 | MHAL_P6 | 2 | 1 | 2 | 3 | 2 | 5 |
| 67 | MHAL_P9 | 1 | 1 | 2 | 1 | 1.25 | 6 |
| 68 | MHSC_A2 | 2 | 1 | 2 | 4 | 2.25 | 6 |
| 69 | MHSC_AI | 2 | 1 | 2 | 2 | 1.75 | 6 |
| 70 | BLC_10 | 2 | 2 | 3 | 4 | 2.75 | 2 |
| 71 | PT_1 | 1 | 2 | 2 | 2 | 1.75 | 6 |
| 72 | PT_41 | 2 | 2 | 2 | 4 | 2.5 | 3 |
| 73 | ORANP_2 | 1 | 1 | 2 | 2 | 1.5 | 7 |
| 74 | ORANP_3 | 2 | 1 | 2 | 2 | 1.75 | 6 |
| 75 | ORANP_6 | 1 | 1 | 3 | 2 | 1.75 | 6 |

| | | | | | | | |
|---|---|---|---|---|---|---|---|
| 76 | ORANR_3 | 1 | 1 | 1 | 1 | 1 | 9 |
| 77 | ORANR_4 | 1 | 1 | 2 | 2 | 1.5 | 7 |
| 78 | ORJEK_I | 1 | 1 | 2 | 2 | 1.5 | 7 |
| 79 | ORPB_15 | 2 | 3 | 3 | 2 | 2.5 | 3 |
| 80 | ORPB_17 | 3 | 1 | 2 | 3 | 2.25 | 4 |
| 81 | ORPB_18 | 2 | 2 | 3 | 4 | 2.75 | 2 |
| 82 | ORPB_20 | 2 | 1 | 3 | 2 | 2 | 5 |
| 83 | ORPB_21 | 2 | 1 | 2 | 2 | 1.75 | 6 |
| 84 | ORPB_9 | 1 | 1 | 2 | 2 | 1.5 | 7 |
| 85 | ORPLM_1 | 2 | 2 | 2 | 3 | 2.25 | 4 |
| 86 | TNT_10 | 2 | 3 | 3 | 3 | 2.75 | 2 |
| 87 | TNT_12 | 1 | 2 | 2 | 3 | 2 | 5 |
| 88 | TNT_13 | 2 | 1 | 2 | 1 | 1.5 | 7 |
| 89 | TNT_14 | 2 | 3 | 3 | 1 | 2.25 | 4 |
| 90 | TNT_15 | 2 | 3 | 3 | 4 | 3 | 1 |
| 91 | TNT_16 | 1 | 2 | 3 | 3 | 2.25 | 4 |
| 92 | TNT_17 | 1 | 1 | 2 | 1 | 1.25 | 8 |
| 93 | TNT_18 | 3 | 1 | 2 | 3 | 2.25 | 4 |
| 94 | TNT_2 | 3 | 2 | 2 | 3 | 2.5 | 3 |
| 95 | TNT_4 | 3 | 1 | 3 | 2 | 2.25 | 4 |
| 96 | TNT_5 | 1 | 1 | 2 | 3 | 1.75 | 6 |
| 97 | TNT_7 | 2 | 3 | 3 | 4 | 3 | 1 |
| 98 | TNT_8 | 2 | 2 | 2 | 1 | 1.75 | 6 |
| 99 | UP_1 | 1 | 1 | 2 | 2 | 1.5 | 7 |
| 100 | UP_C | 2 | 1 | 2 | 3 | 2 | 5 |
| 101 | UP_D | 1 | 1 | 2 | 1 | 1.25 | 8 |
| 102 | UP_E | 1 | 1 | 2 | 2 | 1.5 | 7 |
| 103 | UP_F | 1 | 2 | 2 | 3 | 2 | 5 |
| 104 | UP_K | 1 | 1 | 2 | 2 | 1.5 | 7 |
| 105 | UP_N | 1 | 1 | 2 | 1 | 1.25 | 8 |
| 106 | UP_O | 1 | 1 | 3 | 1 | 1.5 | 7 |

SN- serial number, R_class- resistance class

**Additional Table 2** Genetic information from the 21 co-dominant markers on set of teak genotypes

| SN | Primers | Nucleotide sequences (5’ - 3’)<br>Forward (F) and Reverse (R) | Repeats | Size range (bp) | MAF | GD | Ho | PIC |
|---|---|---|---|---|---|---|---|---|
| 1 | Ver1 | F:CAAAACAAAACCAATAGCCAGAC<br>R: TTTCATCATCATCATCAACATCC | $(GA)_{15}$ | 191-229 | 0.60 | 0.48 | 0.32 | 0.36 |
| 2 | Ver2 | F: AACAACCCCTCCTCTCTTCACTA<br>R:CACTACCACTCATCATCAACACA | $(TC)_{5}$,<br>$(AC)_{5}$ | 236-266 | 0.80 | 0.32 | 0.07 | 0.27 |
| 3 | Ver3 | F: CTTCTGCAACCCTTTTTCAC<br>R: AGCCATATCTTCCTTTCTCT | (GA)20 | 249-279 | 0.54 | 0.50 | 0.29 | 0.37 |
| 4 | Ver4 | F: TTAACGCCAAATCCCAAAG<br>R:CACAAAGAGAACCGACGAG | (TC)10 | 166-170 | 0.67 | 0.44 | 0.20 | 0.34 |
| 5 | Ver5 | F:CGATACCTGCGATGCGAAGC<br>R: CGTTGAATACCCGATGGAGA | (TC)16 | 225-273 | 0.56 | 0.49 | 0.15 | 0.37 |
| 6 | Ver7 | F: AGGTGGGATGTGGTTAGAAGC<br>R: AAATGGTCATCAGTGTCAGAA | (GA)17 | 269-313 | 0.61 | 0.48 | 0.16 | 0.36 |
| 7 | Ver8 | F: AAACCATGACAGAAACGAATC<br>R: TTGGGAATGGGAGGAGAAGT | (GA)16 | 263-291 | 0.54 | 0.50 | 0.25 | 0.37 |
| 8 | Ver9 | F: ATGAAGACAAGCCTGGTAGCC<br>R: GGAAGACTGGGGAATAACACG | (TC)11,<br>(AC)7 | 216-252 | 0.60 | 0.48 | 0.30 | 0.36 |
| 9 | Ver10 | F: CTCGCTTCTTTCCACATT<br>R: ATCATCGCGCATCGTCAA | (AC)10 | 198-222 | 0.68 | 0.43 | 0.08 | 0.34 |
| 10 | Ver11 | F: GCGTCAACCACTTCAACCACCAG<br>R: CCTATTTTCTTCCCCTCCCTTCT | (GA)10 | 204-228 | 0.79 | 0.33 | 0.10 | 0.28 |
| 11 | Ver12 | F: GCTCTCCACCAACCTAAACAA<br>R: AAAACGTCTCACCTTCTCACT | (TC)16 | 198-234 | 0.77 | 0.36 | 0.37 | 0.29 |
| 12 | Ver13 | F: CGCACACCAGTAGCAGTAGCC<br>R: GCCGGAAAAAGAAAAACCAAA | (GA)4 | 129-171 | 0.66 | 0.45 | 0.53 | 0.35 |
| 13 | Ver14 | F: CCGGTAAAAAGGTGTGTCA<br>R: GAGTGGAAGTGCTAATGGA | (TC)4<br>(TC)11 | 217-243 | 0.81 | 0.31 | 0.20 | 0.26 |
| 14 | PAL2 | F:GGCGACCAAGATGATTGAGAG<br>R:CCAGGAAAGTGGAAGACATGAG | | 443 | 0.84 | 0.27 | 0.00 | 0.23 |
| 15 | PAL3 | F:CAGAGCTTGTCAACGACTTCTA<br>R:TTCTGCATCAGCGGGTAAG | | 493 | 0.90 | 0.18 | 0.14 | 0.16 |
| 16 | PAL4 | F:CTGAAGAGCACGGTGAAGAA<br>R:AGCACAGATCGCAGTGAAG | | 489 | 0.69 | 0.43 | 0.00 | 0.34 |
| 17 | CCoAMT1 | F:GGCGACCAAGATGATTGAGAG<br>R:CCAGGAAAGTGGAAGACATGAG | NA | 443 | 0.52 | 0.50 | 0.00 | 0.37 |
| 18 | CCoAMT4 | F:CTGAAGAGCACGGTGAAGAA<br>R:AGCACAGATCGCAGTGAAG | | 489 | 0.71 | 0.41 | 0.00 | 0.33 |
| 19 | CCoAMT5 | F:GCCAATCCCGTTACCAATCA<br>R:GTGTTCTTCACCGTGCTCTT | | 200 | 0.53 | 0.50 | 0.00 | 0.37 |
| 20 | CCR1 | F:ACTGCTCAGAGCTCCAATTC<br>R:TCCTCCCTCACCATCTTGTA | | 607 | 0.74 | 0.38 | 0.52 | 0.31 |
| 21 | Cat | F:CGATTCTCCACTGTCATCCA<br>R:GGAAGTTGTTTCCCACCAAA | | 388-413 | 0.62 | 0.47 | 0.75 | 0.36 |
| | | Average<br>±SD | | | 0.67<br>±0.11 | 0.41<br>±0.09 | 0.21<br>±0.20 | 0.32<br>±0.05 |

MAF- major allele frequency, GD- gene diversity, Ho- observed heterozygosity, PIC- polymorphic information content, SD- standard deviation

**Additional Table 3** Genetic diversity of teak genotypes representing different locality

| States | Accessions | Ne | P% | Ho | He | I |
|---|---|---|---|---|---|---|
| AndhraPradesh (AP) | 31 | 1.67 | 85.71 | 0.37 | 0.36 | 0.52 |
| Arunachal Pradesh (ARP) | 1 | 1.00 | 0.00 | 0.00 | 0.00 | 0.00 |
| Gujrat (GJ) | 1 | 1.00 | 0.00 | 0.00 | 0.00 | 0.00 |
| Keral (KE) | 4 | 1.46 | 71.43 | 0.31 | 0.28 | 0.41 |
| Karnataka (KR) | 17 | 1.65 | 85.71 | 0.37 | 0.36 | 0.52 |
| Maharashtra (MH) | 15 | 1.44 | 76.19 | 0.27 | 0.26 | 0.40 |
| MadhyaPradesh (MP) | 3 | 1.64 | 80.95 | 0.43 | 0.35 | 0.51 |
| Orrisa (OR) | 13 | 1.65 | 85.71 | 0.37 | 0.36 | 0.52 |
| Tamilnadu (TN) | 13 | 1.65 | 85.71 | 0.37 | 0.36 | 0.52 |
| Uttarpradesh (UP) | 8 | 1.64 | 80.95 | 0.37 | 0.35 | 0.50 |
| | Average ±SD | 1.48 ±0.26 | 65.24 ±34.70 | 0.29 ±0.15 | 0.27 ±0.14 | 0.39 ±0.21 |

Ne- number of effective alleles, P%- percentage of polymorphism, Ho- observed heterozygosity, He- Expected heterozygosity/Nei's gene diversity (1972), I- Shannon's information index, SD- standard deviation

**Additional Table 4** Frequencies of the accessions representing different teak populations in the germplasm bank at different resistance class against the pest *E. macheralis*

| Populations | N | $H_o$ | Resistance classes (From Highest (1) to Lowest (9)) | | | | | | | | |
|---|---|---|---|---|---|---|---|---|---|---|---|
| | | | 1 | 2 | 3 | 4 | 5 | 6 | 7 | 8 | 9 |
| AndhraPradesh (AP) | 31 | 0.37 | | | 0.16 | **0.23** | | 0.13 | 0.29 | 0.13 | 0.06 |
| Arunachal Pradesh (ARP) | 1 | 0 | | | | **1.00** | | | | | |
| Gujrat (GJ) | 1 | 0 | | **1.00** | | | | | | | |
| Keral (KE) | 4 | 0.31 | | | 0.50 | | | 0.50 | | | |
| Karnataka (KR) | 17 | 0.37 | | | | 0.12 | | 0.12 | 0.18 | **0.41** | 0.18 |
| Maharashtra (MH) | 15 | 0.27 | | 0.07 | | 0.27 | | 0.13 | **0.40** | 0.07 | 0.07 |
| MadhyaPradesh (MP) | 3 | 0.43 | | 0.33 | 0.33 | | | | 0.33 | | |
| Orrisa (OR) | 13 | 0.37 | | 0.08 | 0.08 | 0.15 | 0.08 | 0.08 | 0.23 | **0.31** | |
| Tamilnadu (TN) | 13 | 0.37 | 0.15 | 0.08 | 0.08 | **0.31** | | 0.08 | 0.15 | 0.08 | 0.08 |
| Uttarpradesh (UP) | 8 | 0.37 | | | | | | 0.25 | | **0.50** | 0.25 |

N- Number of accessions from the population, $H_o$- Observed Heterozygosity

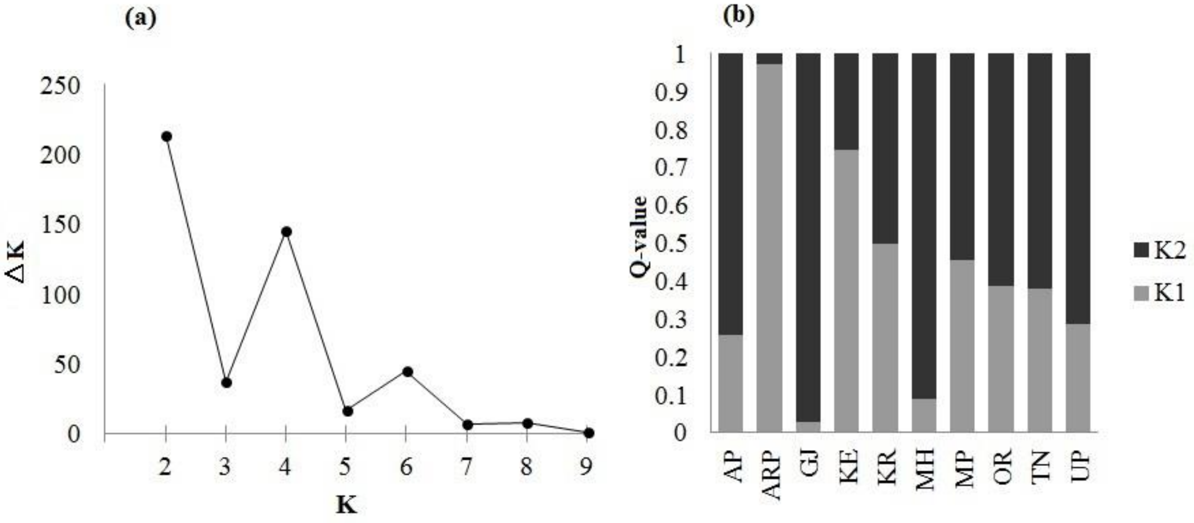


**Additional Figure 1** With very low values for delta K indicating very low genetic structure, STRUCTURE resulted K=2 (a) as the most suitable cryptic number, indicating admixed and unstructured set of teak genotypes (b). Abbreviated name of the states (source locations of teak genotypes) as given in Additional table 3.

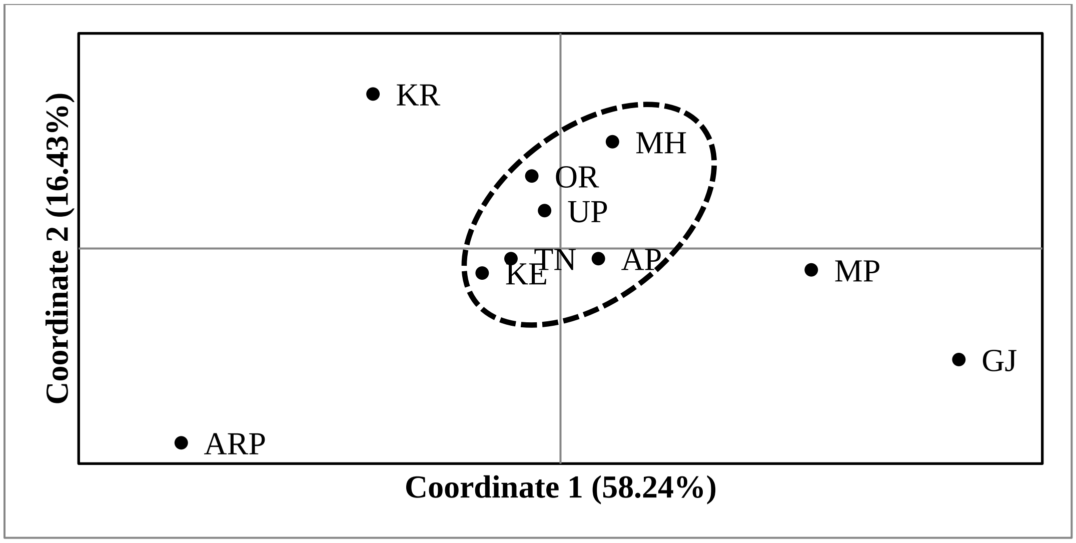


**Additional Figure 2** The PCoA resulted only one clear grouping with admixing of teak populations from north (UP), central (MH, OR) and south (AP, KE and TN) India, covering 74.67% of variation through two axes. Abbreviated name of the states (source locations of teak genotypes) as given in Additional table 3.
